# Estimation of genomic prediction accuracy from reference populations with varying degrees of relationship

**DOI:** 10.1101/119164

**Authors:** S. Hong Lee, Sam Clark, Julius H.J. van der Werf

**Affiliations:** School of Environmental and Rural Science, University of New England, NSW 2351, Australia

**Keywords:** Genomic prediction, Prediction accuracy, Effective number of chromosome segments, Genomic prediction design, varying degrees of relationship

## Abstract

Genomic prediction is emerging in a wide range of fields including animal and plant breeding, risk prediction in human precision medicine and forensic. It is desirable to establish a theoretical framework for genomic prediction accuracy when the reference data consists of information sources with varying degrees of relationship to the target individuals. A reference set can contain both close and distant relatives as well as ‘unrelated’ individuals from the wider population in the genomic prediction. The various sources of information were modeled as different populations with different effective population sizes (*N*_*e*_). Both the effective number of chromosome segments (*M*_*e*_) and *N*_*e*_ are considered to be a function of the data used for prediction. We validate our theory with analyses of simulated as well as real data, and illustrate that the variation in genomic relationships with the target is a predictor of the information content of the reference set. With a similar amount of data available for each source, we show that close relatives can have a substantially larger effect on genomic prediction accuracy than lesser related individuals. We also illustrate that when prediction relies on closer relatives, there is less improvement in prediction accuracy with an increase in training data or marker panel density. We release software that can estimate the expected prediction accuracy and power when combining different reference sources with various degrees of relationship to the target, which is useful when planning genomic prediction (before or after collecting data) in animal, plant and human genetics.

## INTRODUCTION

Genomic prediction of (additive) genetic effects and phenotypes is emerging in a wide range of fields including animal and plant breeding, risk prediction in human medicine and forensics[1-4]. Genomic prediction requires modeling of the association between genome-wide single nucleotide polymorphisms (SNPs) and phenotypes. The success of genomic prediction is measured by its accuracy, i.e. how reliable a future phenotype of target individuals can be predicted.

Genomic prediction requires a reference population of individuals having information on both genotype and phenotype. The accuracy of genomic prediction depends on various parameters, including sample size of the reference and its genetic structure. An important parameter in relation to the latter is the effective size of the population. The effective population size is a predictor of the effective number of chromosome segments that are represented in the population[5-7]. Theoretical predictions have usually considered a homogeneous population of individuals that are essentially unrelated. However, in most practical applications, the reference population used for genomic predictions possibly consists of many sub-groups with individuals having a variety of relatedness to the target individual, e.g. direct relatives, more distant relatives, and individuals that are considered unrelated. It is relevant to assess the contribution of these various sources to prediction accuracy before actually conducting an experiment.

A number of studies have shown that genomic predictions are more accurate if the genomic relationship between the proband and the reference population is higher, both in humans[8-11] and in other species[12-14]. Habier et al (2013)[15] distinguished between three types of information in genomic prediction; linkage disequilibrium, additive-genetic relationships and co-segregation of QTL predicted from markers genotypes with a pedigree. They argued that it would be useful to understand how these sources contribute to the accuracy of genomic predictions, especially when designing reference populations for breeding programs. They show these contributions via simulated examples but did not provide methods that allow simple predictions for their contribution to accuracy. Pszczola et al. (2012)[16] showed that the relationship between the reference population and the proband should be maximized to achieve an optimal design using a simulation study. However, they also did not attempt to derive the expected prediction accuracy from an optimal design in advance. Hayes et al. (2009)[17] considered the influence of direct relatives on genomic prediction. They followed the same approach as the general theory, i.e. by considering the number of independently segregating chromosome segments within families. They showed the accuracy of genomic prediction from varying sizes of the first and second degree of relatives, but did not consider the information from combined sources[18]. It should also be noted that those studies that derived genomic prediction accuracy from theory using effective number of chromosome segments (*M*_*e*_)[5, 6, 19-21], did not consider the correlation between relatedness at different chromosomes, therefore overestimating *M*_*e*_ and underestimating whole-genome prediction accuracy[7].

Wientjes et al (2016)[22] proposed a simple selection index approach to combine information from different populations. They considered a genetic correlation between genetic effects expressed in different populations. We propose to use the same approach to combine different sources of information from different subsets within a population, where the different subsets have a different degree of relationship with the target individual. To predict the accuracy, we derive the number of effective chromosome segments from a hypothetical *N*_*e*_ associated with each subset, and we show that that combining such subsets using selection index theory gives the same result as using a prediction from an *M*_*e*_ derived from the variation in genomic relationships between the overall reference data and the target. Prediction accuracy is derived from variation in genomic relationship rather than the variation in genomic relationship as a deviation for the expected relationship among members of the reference set, as was proposed by Goddard et al (2011) and also applied by Wientjes et al (2016). This approach leads to a theoretical concept useful for assessing the accuracy of genomic predictions in advance, and we illustrate this with examples based on real data.

## MATERIALS AND METHODS

### Predicting genomic selection accuracy

The accuracy of genomic breeding values (GBV) or (genomic profile score in the context of human risk prediction[23]) based on genome-wide SNP genotypes can be predicted from theory[5-7, 24], assuming that prediction is based on a reference population with phenotypes and genotypes for the same genome-wide SNPs that are linked to quantitative trait loci (QTL). The accuracy depends on i) the proportion of genetic variance at QTL captured by markers and ii) the accuracy of estimating marker effects. The proportion of genetic variance at QTL captured by markers (b) depends on linkage disequilibrium (LD) between markers and QTL, which in turn depends on the number of markers (*M*) and the number of ‘effective chromosome segments’ (*M*_*e*_)[5], that is

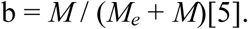

Various forms of prediction of *M*_*e*_ have been presented[5, 6, 21] that were however inconsistent to each other, and without considering the correlation between chromosomes. Recently, we presented a prediction formula with the form[7]

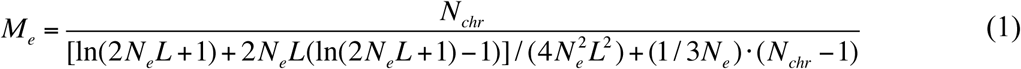

where *N*_*e*_ = effective population size; *L* = average chromosome length; *N*_*chr*_ = number of chromosomes. This formula accounts for mutation, and that without considering mutation should be referred to equation (10) in Lee et al. (2017)[7]. If *N*_*chr*_ = 30 and *L* = 1, Eq. (1) can be approximately simplified as

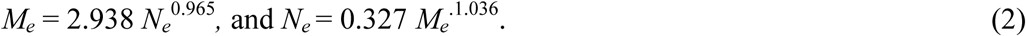

The accuracy of the genomic prediction of a phenotype can be written as[5]

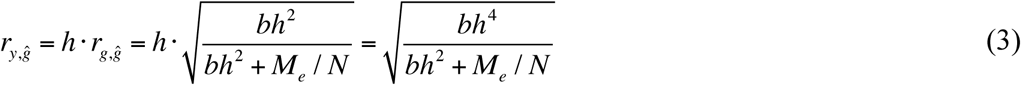

where 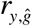 is the correlation coefficient between the true phenotypes (*y*) and estimated GBV, *h*^*2*^ is the heritability of the trait, and *N* is the number of phenotypic observations. Other measures for genomic prediction accuracy, particularly for human risk prediction, such as the area under the receiver operating characteristic curve (AUC) or odds ratio of case-control status contrasting the higher or lower risk group are described elsewhere[7].

### *M*_*e*_ and genomic relationship

After collecting genotypic information of the reference data and the target individual, it is possible to obtain an empirical *M*_*e*_ from a genomic relationship matrix (GRM). The elements in the GRM are 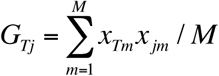 where *x*_*Tm*_ and *x*_*jm*_ are the standardised genotype coefficients (mean 0 and variance 1) for the target individual (*T*) and *j*th individual in the reference data at the *m*th locus. It is possible to construct a GRM for each locus, and the elements in the GRM at the *m*th locus are *G*_*Tj* (*m*)_ = *x*_*Tm*_*x*_*jm*_. Then, the variance of the mean of *G*_*Tj* (*m*)_ across all *M*_*i*_ SNPs in a single chromosome is

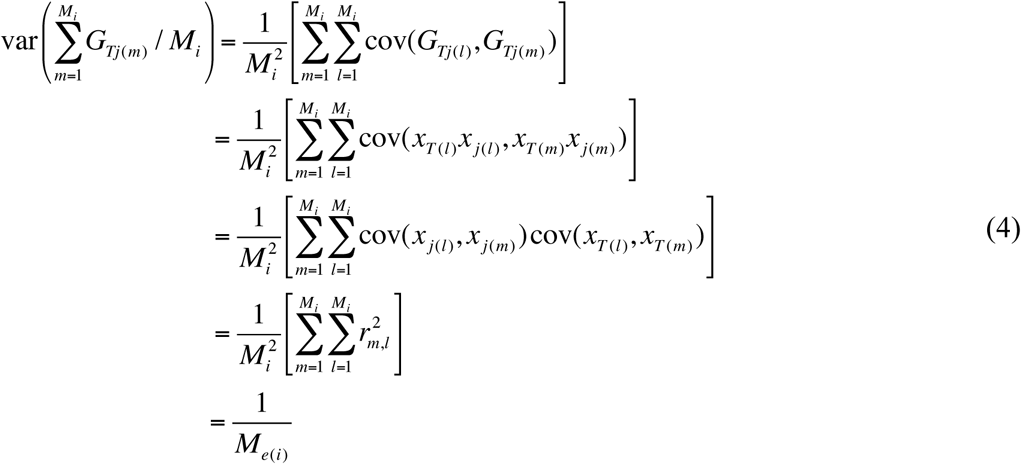

where cov(*x*_*j* (*l*)_, *x*_*j* (*m*)_) = cov(*x*_*T* (*l*)_, *x*_*T* (*m*)_) = *r*_*m*,*l*_, which is a correlation between the *m*^th^ and *l*^th^ SNP-genotype, because of var(*x*) = 1 and mean(*x*) = 0, i.e. the genotype coefficients are standardized in the population. The term *M*_*e(i)*_ is the effective number of chromosome segments for the *i*th chromosome, and it is calculated from the inverse of the average of the squared correlation between the *M*_*i*_ SNPs[6, 7]. When considering multiple chromosomes, the covariance of the pairwise relationship between two chromosomes is not negligible[7]. Assuming equal length and number of SNPs for *N*_*chr*_ chromosomes, *M*_*e*_ for the whole genome can be derived from the variance in the relationship across the whole genome as

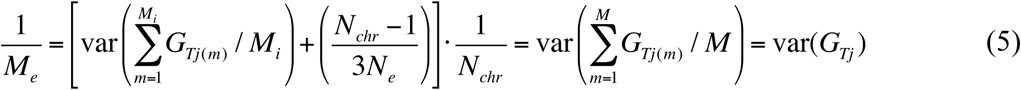

where *N*_*chr*_ is the number of chromosomes. This expression is equivalent to (1) when using Sved (1971)[25] for deriving the expected squared correlation between genotypes, which is dependent on *N*_*e*_. An empirical *M*_*e*_ can be derived from the GRM as the variance in relationships between a target individual *T* and *N* individuals in the reference population. Goddard et al. (2011)^5^ suggested their theoretical derivation had to assume a homogeneous population of individuals that are essentially unrelated. However, Eq. (4) show that the assumption about unrelated individuals is not necessary so that any random samples from the population can be used, irrespective of whether they are highly related or not (see Results).

### Effective population size in a reference data set

One of critical parameters to determine the accuracy of genomic prediction is the effective population size (*N*_*e*_). It is not very common to represent a reference population by a single value of *N*_*e*_ when it consists of several cohorts of individuals with different relationships to the target individual. Wientjes et al. (2016)[22] used a single value for *M*_*e*_ representing a reference set consisting of two populations. Here, we generalized that concept for any number of subsets of the reference population based on the relationship between *N*_*e*_, *M*_*e*_ and var(*G*_*T\**_), leading to a certain value for *M*_*e*_ and *N*_*e*_ for a reference population consisting of several cohorts. For any subset of the reference data set, there are realized relationships with the target sample. From Eq. (5), a value of *M*_*e*_, which is the inverse of the variance of the genomic relationships between the target and the reference sample, can be assigned to the reference data. Then, a single value of *N*_*e*_, which is a function of *M*_*e*_ from Eq. (1) or (2), can be obtained for the reference data. The effective population size of the reference set is therefore a parameter specific to the data used and it describes genomic diversity of the reference individuals used for prediction of genomic breeding value relative to the target individual which breeding value is being predicted. This *N*_*e*_ value can be smaller than the effective size of the population from which the reference individuals were sampled, but it can also be larger, depending on whether the reference individuals chosen were closer or more distantly related to the reference set.

Based on this concept of *N*_*e*_, reflecting information content of the reference sample in relation to the target sample, we consider three information sources consisting of i) close relatives of the proband, e.g. *N*_*e*_ = 10, ii) distant relatives or individuals from the local area of the proband, e.g. *N*_*e*_ = 100 and iii) a wider population sample of individuals that are not related to the proband, e.g. *N*_*e*_ = 1,000.

The GBV can be estimated based on each of these information sources, and the accuracy of the estimation can be calculated as above, e.g. 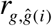 from Eq. (3) where *i* represents the *i*th information source. It is also possible to estimate GBV based on combined data of all three information sources. Assuming a random sample from the same population for each source, the accuracy of the GBV based on the combined data set can then be calculated using standard selection index theory as

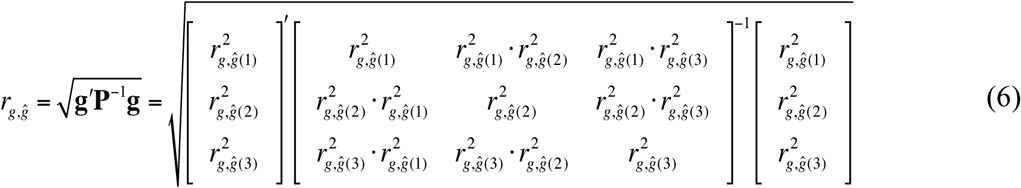

where **g** is the vector with covariances between each of the GBV and the true breeding value, and **P** is the variance-covariance matrix of the set of GBV. The accuracy of the GBV based on the combined data set can also be estimated based on the weighted *M*_*e*_ from the three information sources. Assuming a random sample from the same population for each source, the weighted *M*_*e*_ can be obtained as

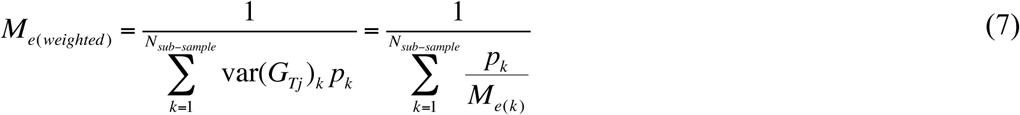

where *p*_*k*_ is the proportion of the sample size over the total sample for each information source. The accuracy of the GBV based on the weighted *M*_*e*_ is identical with that using standard selection index theory above (Eq. (6)).

Following Wientjes et al. (2016)[22] we can further generalize for a case where genetic correlations among multiple reference populations and those between reference populations and the target are not one. Equation (6) can be generalized as

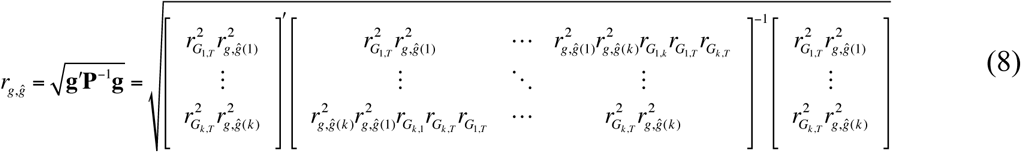

where 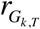 is the genetic correlation between the *k*^th^ reference population and the target set, and similarly, 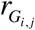 is the genetic correlation between the *i*^th^ and *j*^th^ reference population (*i* = *j* = 1 ∼ k).

equation (8) can be used when multiple reference populations have both quantitative traits and diseases. If the k^th^ reference population is measured for binary response (disease or disorder), the reliability term, 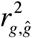, in Eq. (8) can be replaced with

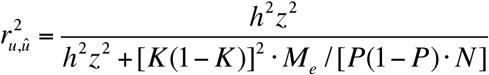

where *u* is a genetic profile score on the 0,1 disease scale[26, 27], *K* is the population prevalence for the disease, *P* is the proportion of cases in the total sample *N* of cases and controls, and *z* is the density at the threshold on the normal distribution in the liability threshold model.

When the target data set is measured for a binary response (e.g. diseases), the AUC or odds ratio of case-control status contrasting the higher or lower risk group can be also estimated[7]. These measures for genomic prediction accuracy with multiple heterogeneous reference populations can be obtained using MTG2, publicly available software (https://sites.google.com/site/honglee0707/mtg2).

### Power of genomic prediction

The power depends on the sample size in the target data set and the overall reliability of the genomic prediction (Eq. (8)).

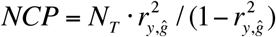

where NCP is the non-centrality parameters, N_T_ is the sample size in the target data and 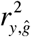 is the coefficient of determination of the predictor. Then, the power can be written as

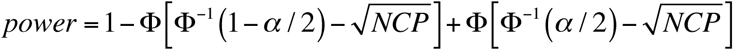

where Φ is the cumulative standard normal function and α is the significance level. When using the odds ratio of case-control status contrasting the higher or lower risk group, the non-centrality parameters can be derived as[28]

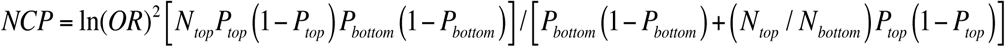

where *N*_*top*_ and *N*_*bottom*_ is the number of individuals in the top and bottom percentile, and *P*_*top*_ and *P*_*bottom*_ is the proportion of cases in each group. The power can be estimated as above.

### Simulation

In a simulation, a stochastic gene-dropping method [29, 30] was used to simulate 4,000 SNPs for each of 30 chromosomes, each of length *L*=1 Morgan with *N*_*e*_ = 50, 500 and 1000 for 50, 500 and 1000 generations, respectively. Recombination and mutations were modelled according to the genetic distance between SNPs and the mutation rate of 1e-08 per site per generation[31]. In the final generation, we constructed a genomic relationship matrix for a random set of 3000 individuals. Among the 3000 individuals, we randomly selected 1000 individuals as target data and 2000 individual as reference data, and estimated variance of the genomic relationships between the target and reference data to validate Eq. (1), (2), (4) and (5).

### Evaluation of the formulas

For each of the three information sources contributing to genomic prediction we varied values for *N*_*e*_, sample size in reference data and marker density. We compared the expected accuracy of GBV from the sample of *N*_*e*_ =1000 with predictions that additionally included information from the sample of *N*_*e*_ =100 and *N*_*e*_ =10. The total number in the reference population was kept equal between the comparisons.

### Real data analysis

We used publicly available data from the Framingham heart study (phs000007.v26.p10.c1)[32]. There were 6950 individuals genotyped for 500,568 SNPs. Stringent quality control for genotype data and phenotype adjustment for confounders were applied to the data (the details can be found in Lee et al. (2016)[7]). The quality control included SNP call rate > 0.95, individual call rate > 0.95, HWE p-value > 0.0001, MAF > 0.01 and individual population outliers < 6 SD from the first and second principal components (PC). After QC, 6920 individuals and 389,265 SNPs remained. Among them, 4243 individuals were phenotyped for height and body mass index (BMI).

We made three different information sources to form the reference data that were tested in 100 replicated analyses (Table 1). Initially, we randomly selected 800 individuals out of 4243 phenotyped individuals as a target data set. For reference data set #1, we selected 50% of individuals that were highly related (> relatedness of 0.3) to the 800 target individuals (*N*_*1*_ = 617 ± 19). For reference data set #2, we selected 80% of moderately related individuals (> relatedness of 0.1) of the 800 target individuals (*N*_*2*_ = 1254 ± 30). For reference data set #3, we took the rest of the individuals that were not selected for reference data set #1 and #2 (*N*_*3*_ = 1572 ± 33). There was no overlap sample between target data set and reference data sets #1, #2 and #3.

**Table 1.**
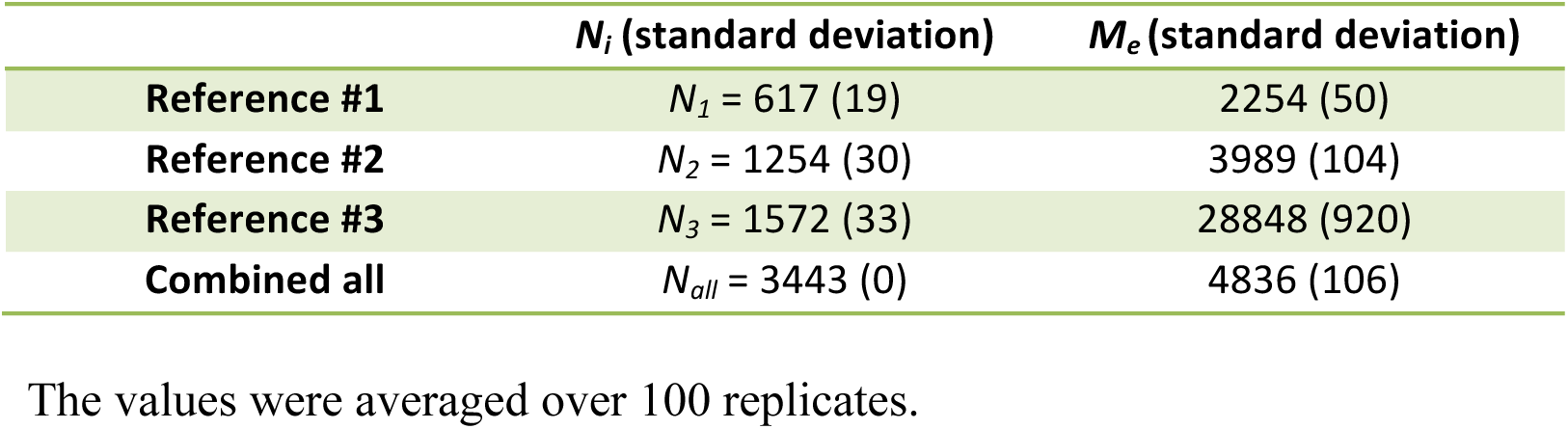
The sample size (*N*) and empirically observed *M*_*e*_ in each of three different reference data sets and combined data set in the Framingham data analysis.

| | $N_i$ (standard deviation) | $M_e$ (standard deviation) |
| --- | --- | --- |
| **Reference #1** | $N_1 = 617$ (19) | 2254 (50) |
| **Reference #2** | $N_2 = 1254$ (30) | 3989 (104) |
| **Reference #3** | $N_3 = 1572$ (33) | 28848 (920) |
| **Combined all** | $N_{all} = 3443$ (0) | 4836 (106) |

The values were averaged over 100 replicates.

Using the real genotype data, the genomic relationships between the reference and target sample were constructed. Empirical *M*_*e*_ was estimated from equation (5) for reference #1, 2 and 3, and that for combined data. We took a median rather than mean because the distribution of variance of the genomic relationship between target and reference sample was skewed. The correlation between the true phenotypes (that were not used in the analyses) and estimated GBV in the target data set was estimated for the combined data set, which was used as the genomic prediction accuracy 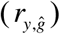. Phenotypes were adjusted for birth year, sex, and the first 10 PCs were used to control non-genetic confounding effects, e.g. population stratification.

## RESULTS

In the simulation study, as shown in Figure 1A, 1B and 1C, the expected (from Eq. (1)) and empirically observed *M*_*e*_ from the simulated genotyped data (using Eq. (5)) are in good agreement, however, they are considerably lower than the expectation from the previous formulas[5, 6, 21], which confirms the result from Lee et al. (2017)[7]. It is noted that Eq. (5) is still valid in the subset with a smaller *N*_*e*_ = 50 that has a significant proportion of high related individuals, indicating that the assumption about unrelated individuals (made in Godard et al. (2011)[5]) can be relaxed. It is shown that whether using a high or low effective population size, the mean and variance of genomic relationships is generally agreed with the expectation from the theory (Supplementary Figures S1 and S2).

**Figure 1.**
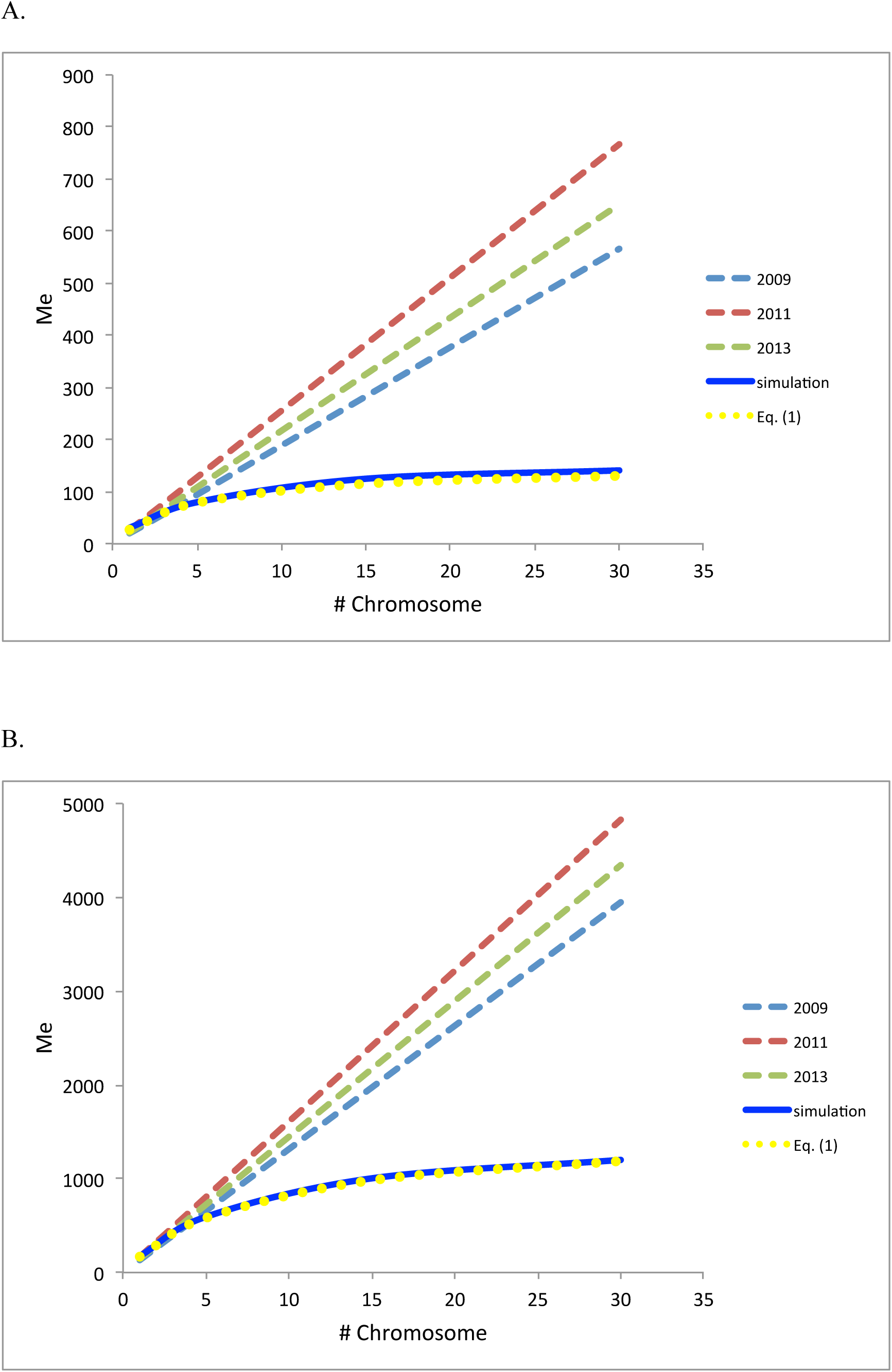
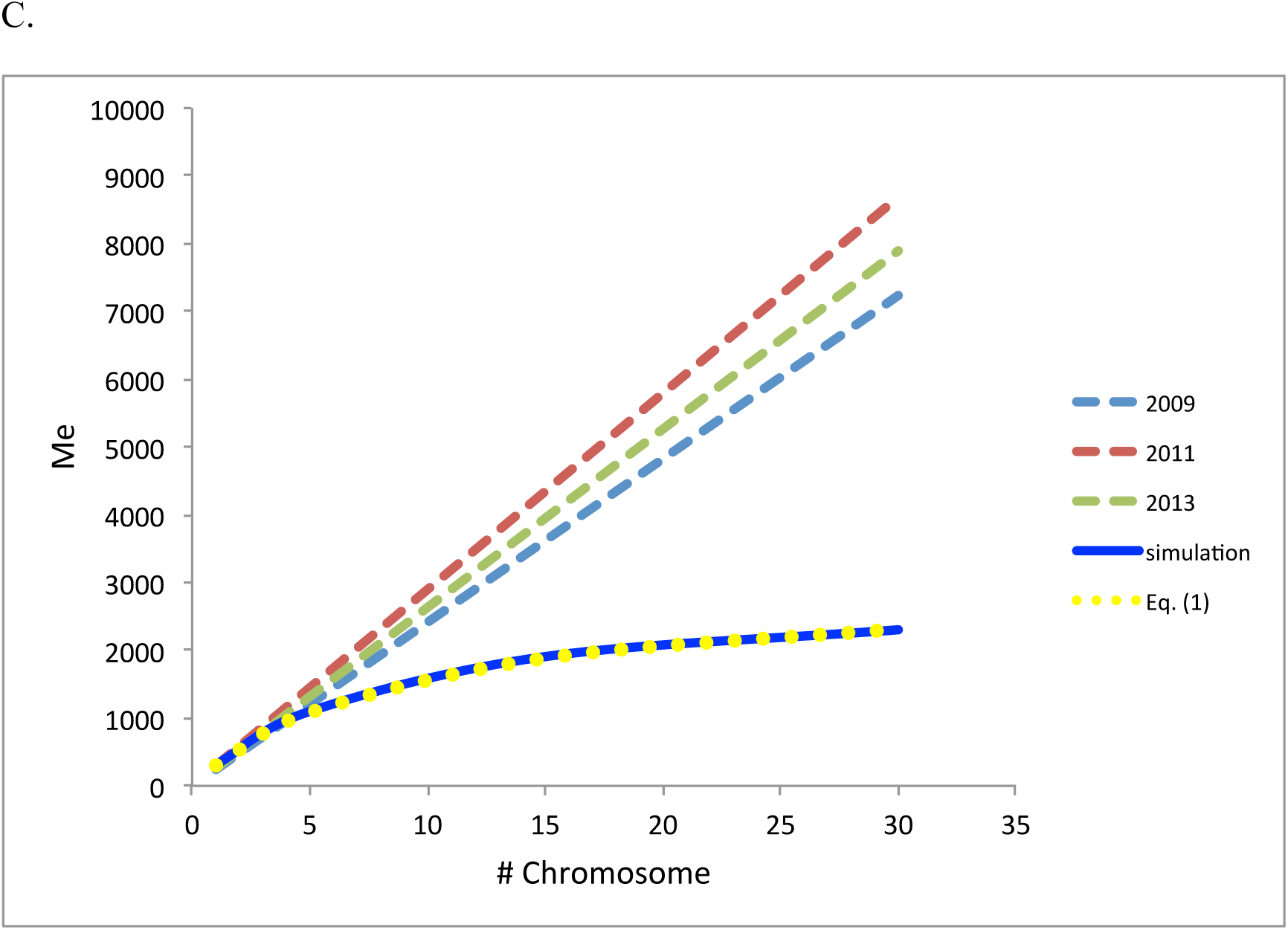
Expected effective number of chromosome segments (*M*_*e*_) from previous studies in 2009[6], 2011[5] and 2013[21] and from Eq. (1) in this study, compared to empirically observed from *M*_*e*_ simulation when varying the number of chromosomes each with 1 Morgan long. Effective population size was used as *N*_*e*_ =50 **(A)**, 500 **(B)** and 1000 **(C)**. This confirms the result from Lee et al. (2017)[7].

In the evaluation of the formulas, we first tested how the prediction accuracy was changed with varying marker density, using formula (1) and (3) and b = *M* / (*M*_*e*_ + *M*) (Figure 2). For *N*_*e*_=10,000, the accuracy gradually increased with marker density, but the slope became flat when using the number of SNPs exceeded 100,000 (Figure 2A).

**Figure 2.**
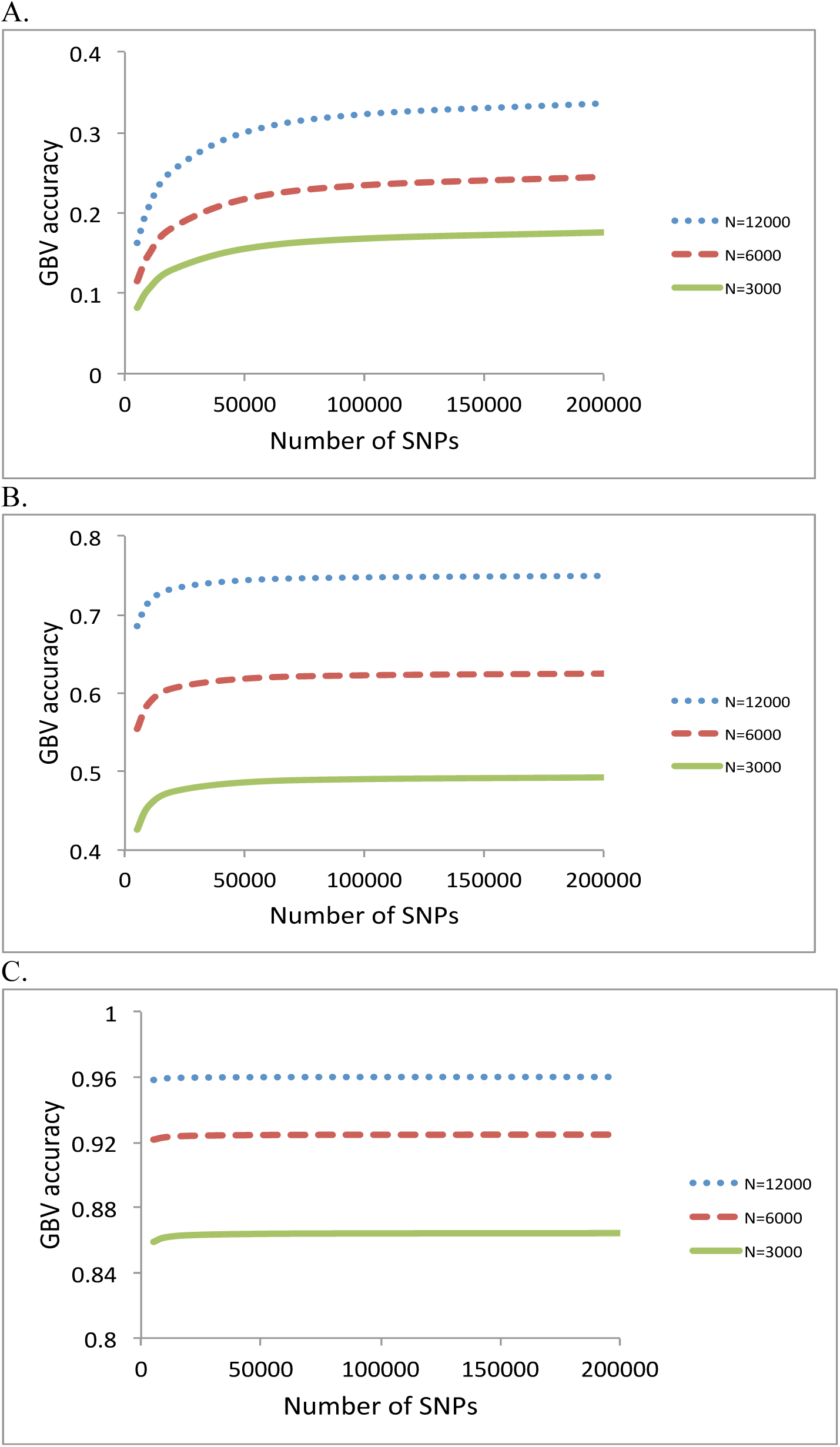
Accuracy of GBV when varying the number of SNPs for *N*_*e*_ = 10,000 **(A)**, 1000 **(B)** and 100 **(C)**. The sample size in the reference data was *N*=12,000, 6000 or 3000. The heritability was 0.25.

For *N*_*e*_=1,000, the accuracy did not increase with marker density as long as the number of SNPs was higher than 50,000 (Figure 2B). For *N*_*e*_=100, there was no improvement of the accuracy if the number of SNPs was more than 10,000 (Figure 2C). This would be expected because the proportion of genetic variance at QTL captured by markers (b = *M* / (*M*_*e*_ + *M*) approached one when the number of SNPs (*M*) was more than 100,000, 50,000 and 10,000 for *N*_*e*_ = 10,000, 1000 and 100, respectively (Figure 3), as *M*_*e*_ was equal to 21,248, 2,313 and 254 for these three cases.

**Figure 3.**
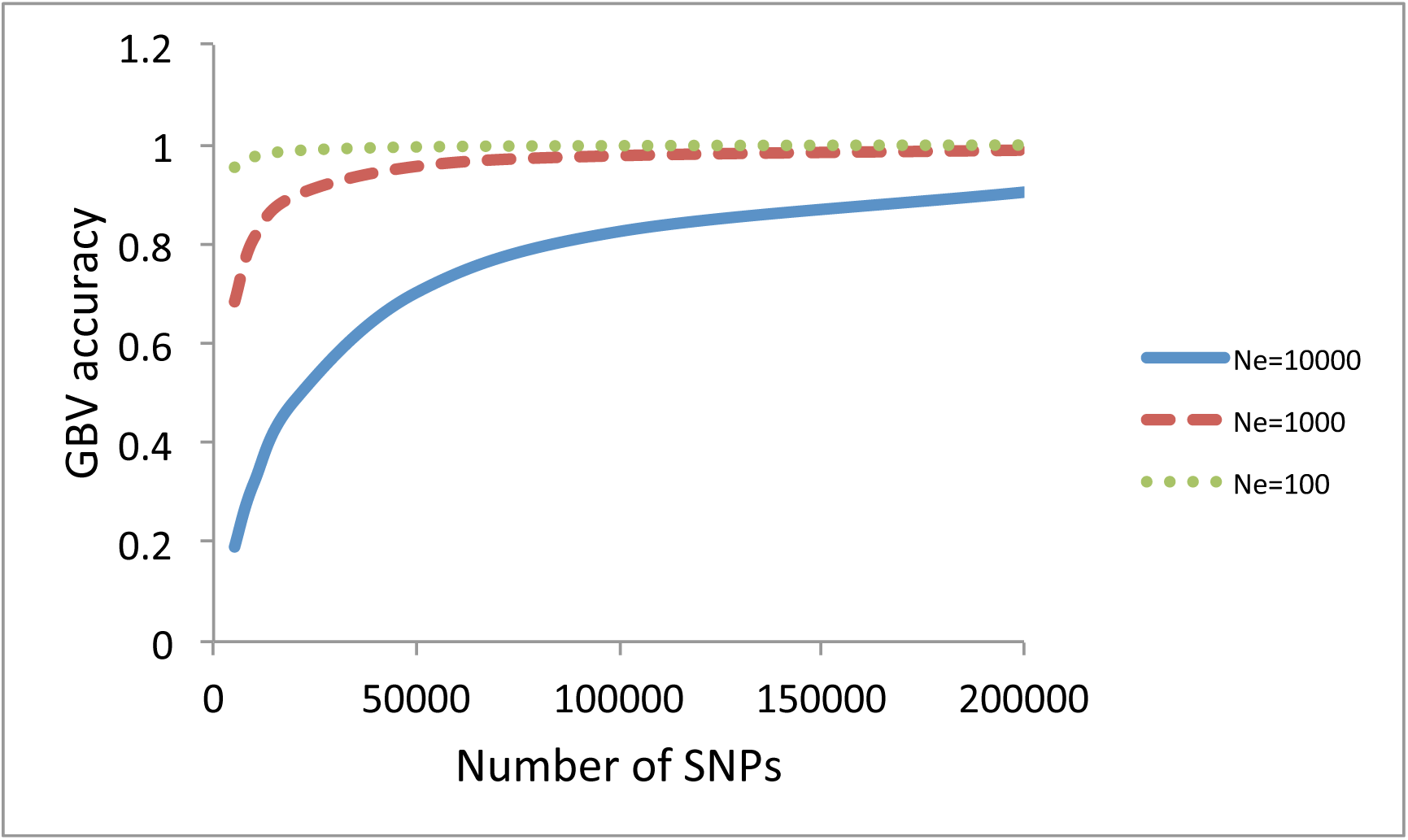
The proportion of genetic variance at QTL captured by markers (b = *M* / (*M*_*e*_ + *M*) when varying the number of SNPs for *N*_*e*_ = 10,000, 1000 and 100.

Next, we quantified the contribution of each information source when varying sample size in the reference data using formula (1), (3) and (5) (Figure 4). It was assumed that the number of SNPs was sufficient to capture most of causal variants (e.g. > 50,000). When adding 100 individuals of *N*_*e*_=100 or *N*_*e*_=10 to the reference sample with *N*_*e*_=1000, the accuracy was slightly or substantially improved (Figure 4A). The improvement was larger when adding more individuals (500) (Figure 4B). Results showed that an information source of a smaller *N*_*e*_ was more important when the samples sizes of each information source were the same. When the total number of reference data was increased, the importance of adding an information source of a smaller *N*_*e*_ was relatively decreased (Figure 4). When heritability was higher, overall accuracy was increased, and the relative contribution from an information source of a smaller *N*_*e*_, i.e. the close relatives, was reduced (Figure 5).

**Figure 4.**
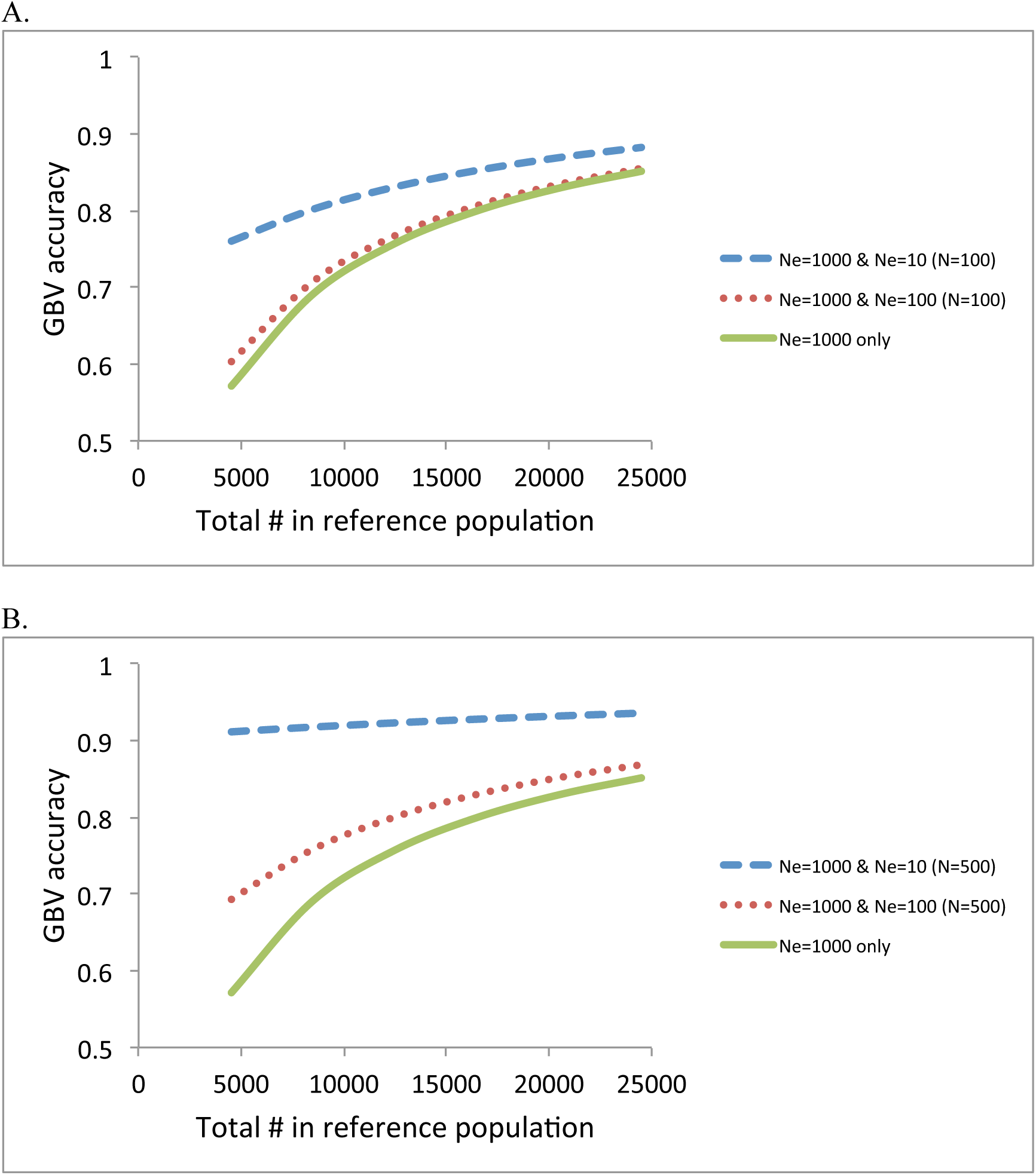
Accuracy of GBV when adding 100 individuals (*N*=100) **(A)** or 500 individuals (*N*=500) **(B)** of *N*_*e*_=100 or *N*_*e*_=10 to the reference population of *N*_*e*_=1000. The heritability was 0.25.

**Figure 5.**
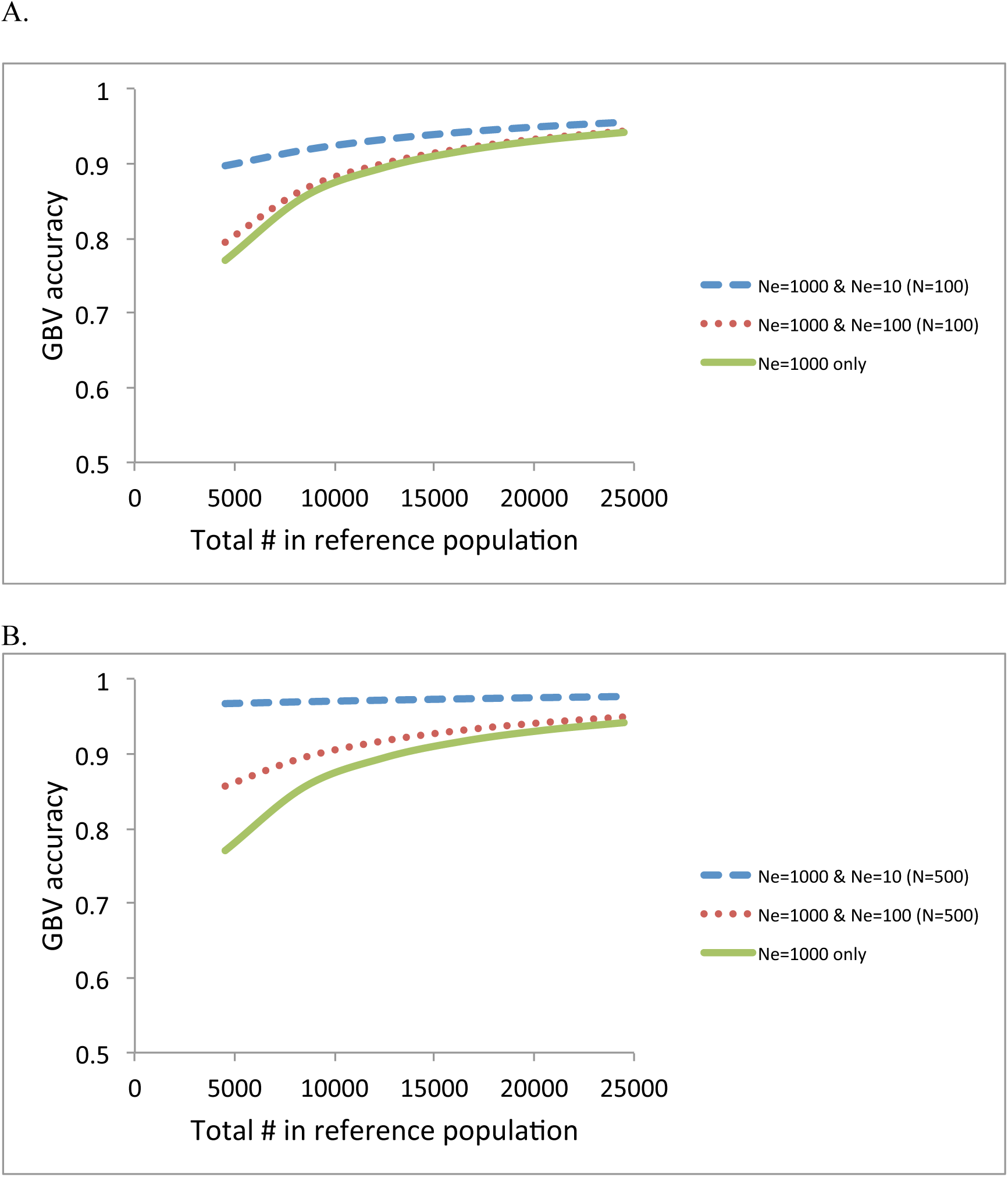
Accuracy of GBV when adding 100 individuals (*N*=100) **(A)** or 500 individuals (*N*=500) **(B)** of *N*_*e*_=100 or *N*_*e*_=10 to the reference population of *N*_*e*_=1000. The heritability was 0.25. The heritability was 0.75.

Figure 6 confirms again that the smaller *N*_*e*_, the better the prediction accuracy when using each information source separately. However, the sample sizes can be also varied across the information sources, as there are generally a lot fewer close relatives than individuals from the wider population. In Figure 6A, the accuracy at a sample size of 100 for *N*_*e*_=10 was 0.73, which was lower than that of a sample size of 1,000 for *N*_*e*_=100 (0.81) or that of a sample size of 20,000 for *N*_*e*_=1,000 (0.83). With a higher heritability, the result is similar in that the 20,000 records in the information source of *N*_*e*_=1000 gave a better accuracy than the 100 records of close relatives (*N*_*e*_=10).

**Figure 6.**
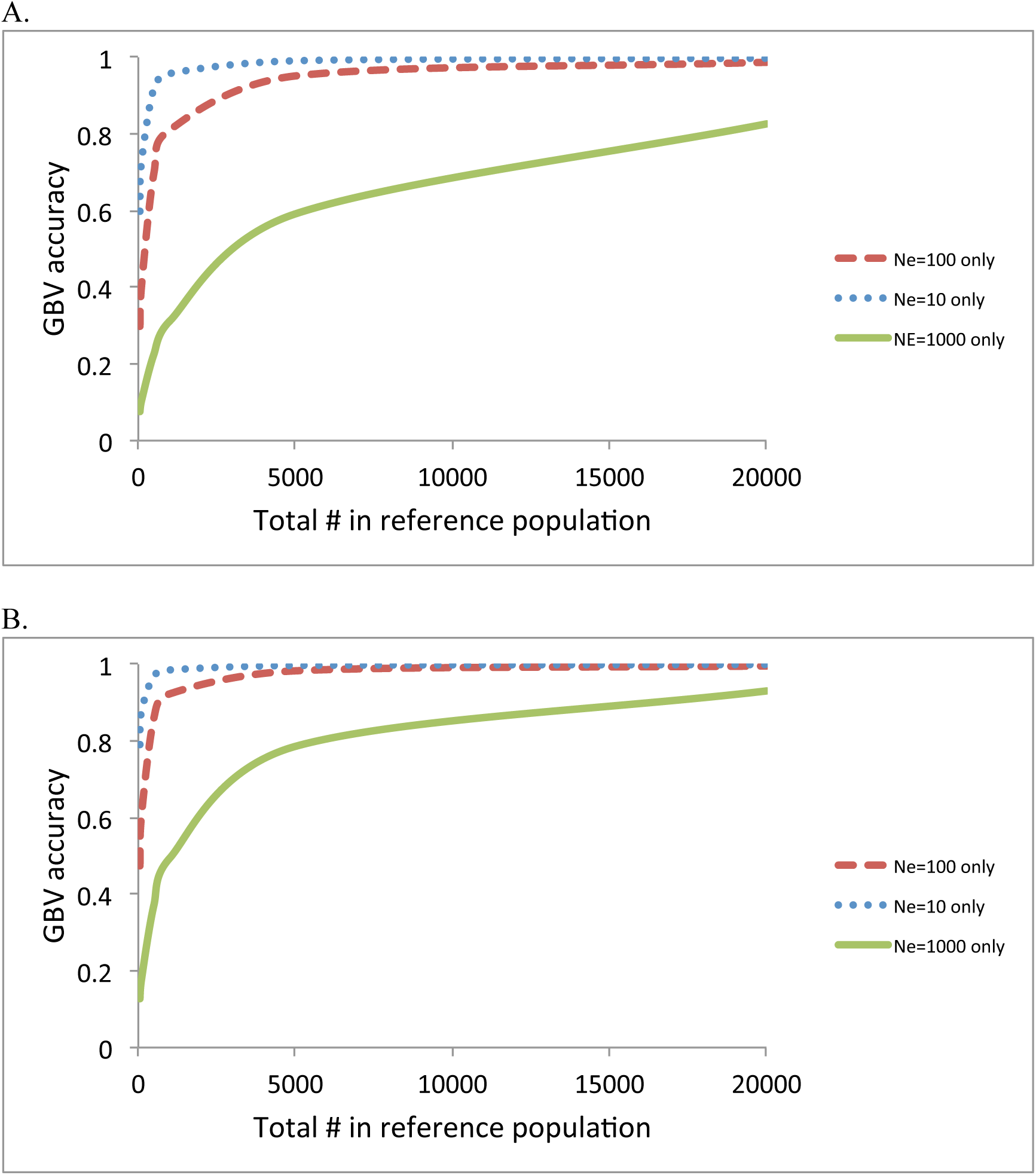
Accuracy of GBV when using *N*_*e*_=1000 only, *N*_*e*_=100 only and *N*_*e*_=10 only with a heritability of 0.25 **(A),** and with a heritability of 0.75 **(B).** For *N*_*e*_=10 only, the accuracy at a sample size of 100 was 0.73 **(A)** and 0.88 **(B).** For *N*_*e*_=100 only, the accuracy at a sample size of 1000 was 0.81 **(A)** and 0.92 **(B).** For *N*_*e*_=1000 only, the accuracy at a sample size of 20,000 was 0.83 **(A)** and 0.93 **(B).**

In real situations, the most common and desirable design may combine all of the information sources to maximize the prediction accuracy. We plotted the accuracy using a composite design consisting of *N*_*e*_=1000 + *N*_*e*_=100 (*N*=500) + *N*_*e*_=10 (*N*=50), compared to that using *N*_*e*_=1000 (Figure 7). The accuracy for a composite design was substantially increased especially when the total number of reference sample is low (Figure 7).

**Figure 7.**
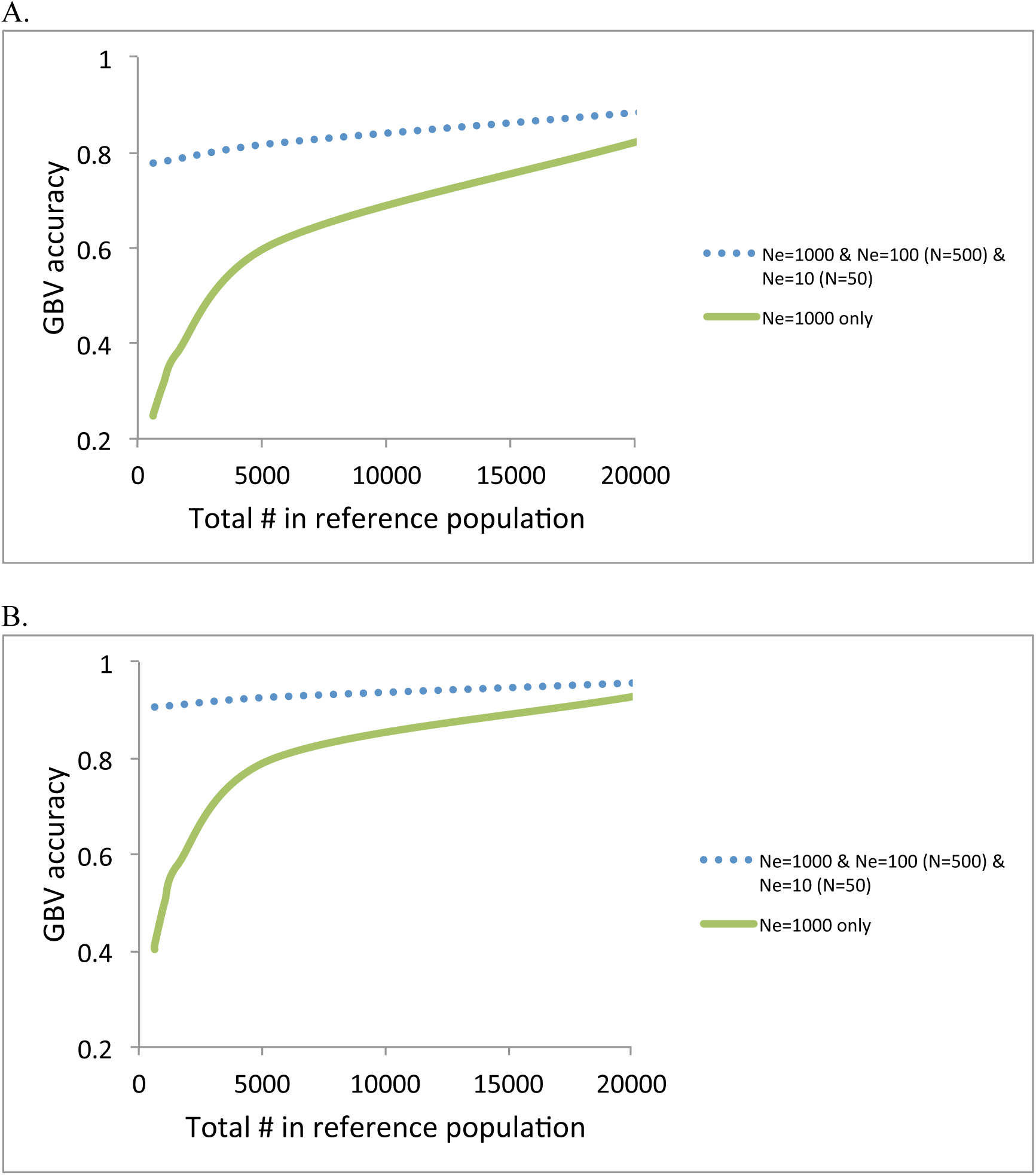
Accuracy of GBV when using a composite design, e.g. *N*_*e*_=1000 + *N*_*e*_=100 (*N*=500) + *N*_*e*_=10 (*N*=50), compared to Ne=1000 only with a heritability of 0.25 **(A)** and with a heritability of 0.75 **(B).**

Figure 8 illustrates the real data analyses. The median of empirically estimated *M*_*e*_ from the inverse of the variance of the genomic relationship (Eq. 5) over 100 replicates was 2254 (SD=50), 3989 (SD=104) and 28848 (SD=920) for reference #1, #2 and #3, respectively (Table 1). Empirically estimated *M*_*e*_ based on the combined data was 4836 (SD=106) while expected *M*_*e*_ was 5309 (SD=88), approximately confirming Eq. (7). The (small) difference between empirical observation and expectation was probably due to skewed distribution of the variance of the genomic relationships.

**Figure 8.**
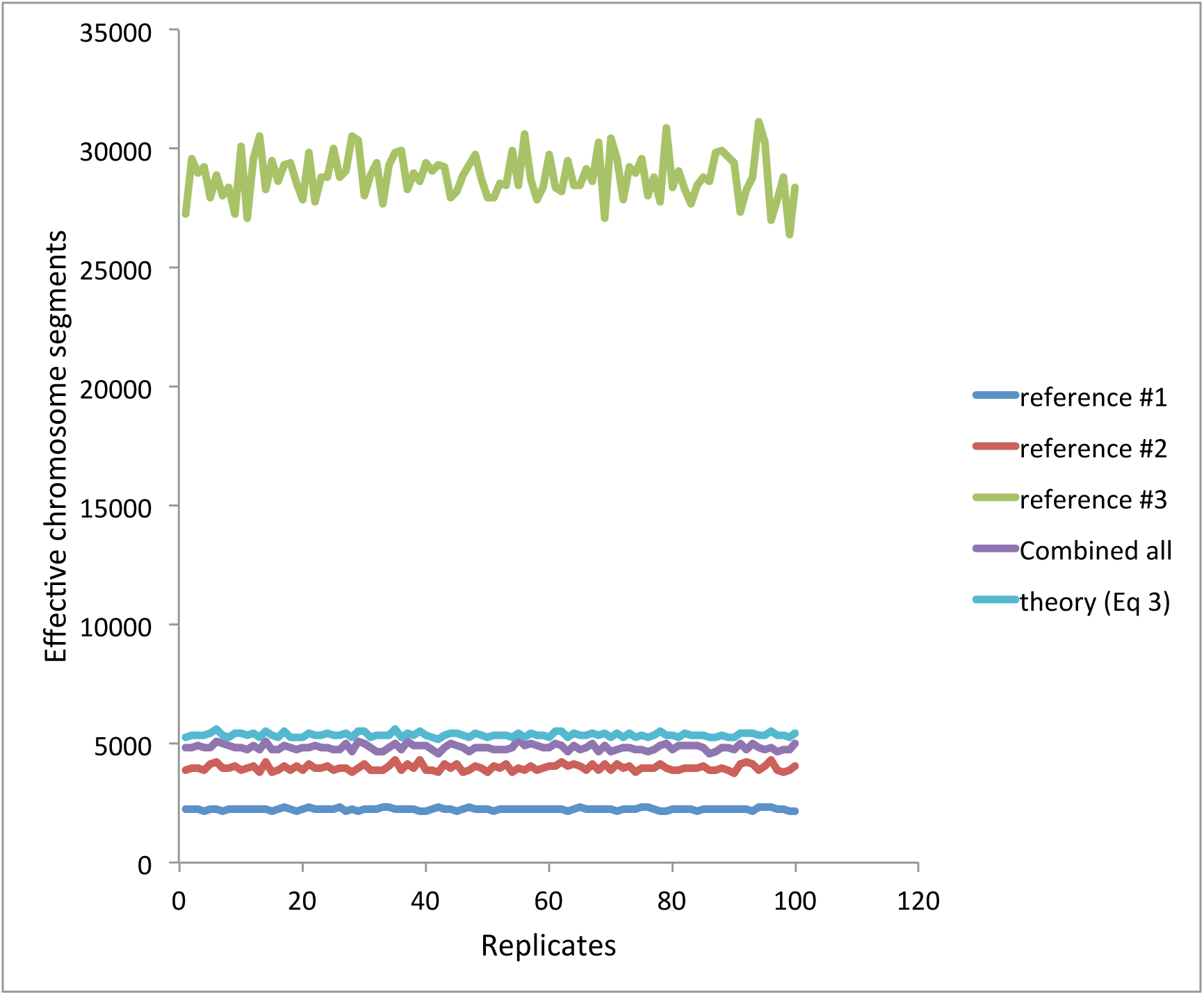
When using Framingham data, empirically estimated *M*_*e*_ based on each of the reference data sets and combined data. Empirically estimated *M*_*e*_ based on combined data is approximately agreed with that from theory (Eq. 5).

Given *M*_*e*_, *N* and *h*^*2*^, the expected accuracy agreed well with the observed accuracy when using Framingham data (Figure 9). The reported heritabilities, *h*^*2*^=0.8 [33-35] for height and *h*^*2*^=0.46[36, 37] for BMI, were used. We also quantified the importance of marker density using the real data. In agreement with Figure 2, the prediction accuracy is not much decreased even with 50,000 SNPs that were randomly selected from 389,265 SNPs (Figure 10).

**Figure 9.**
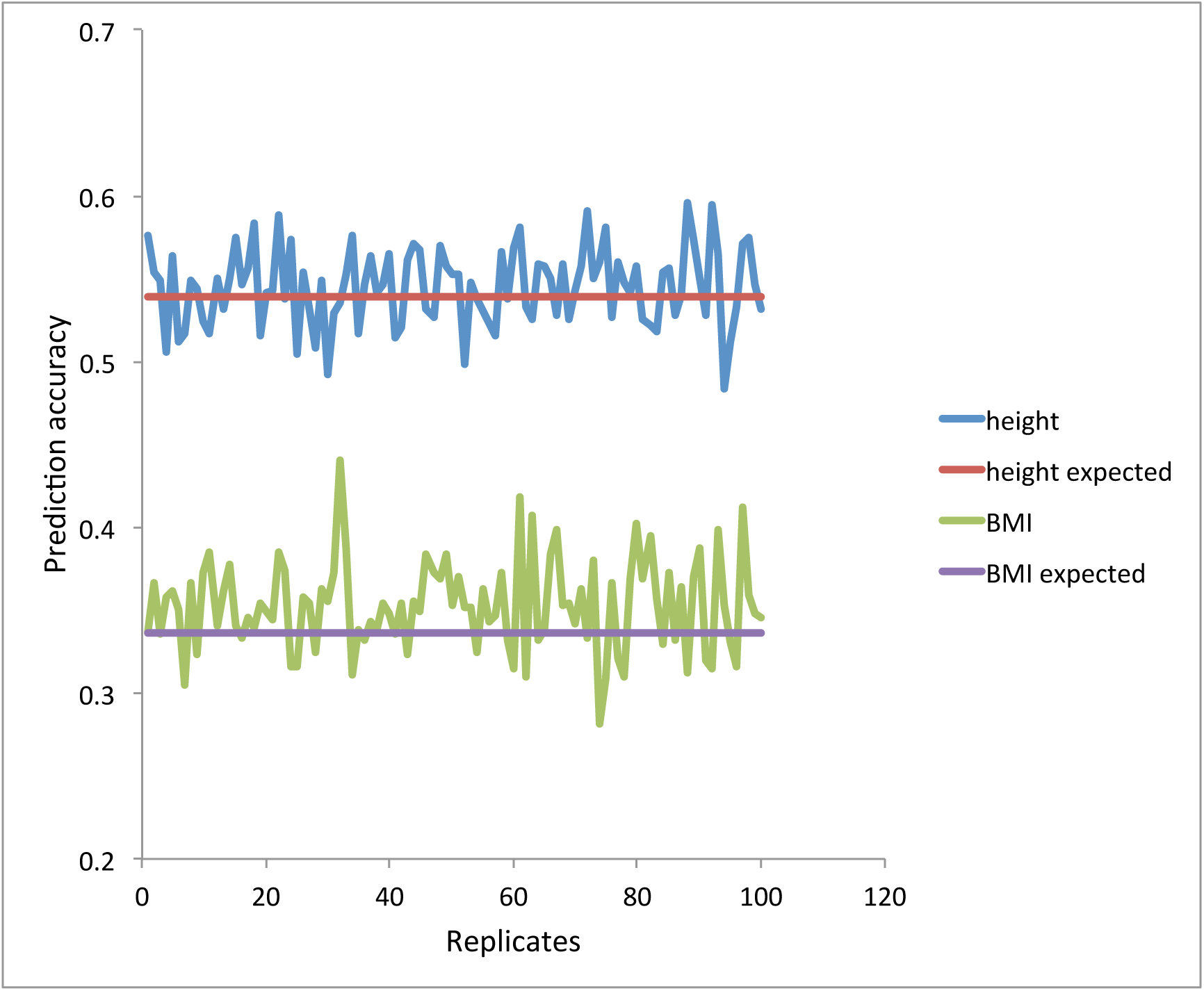
When using Framingham data, observed prediction accuracy and expected prediction accuracy with given *M*_*e*_ and *N* (from Eq. (3)) are agreed well. The reported heritability, *h*^*2*^=0.8 [33-35] for height and *h*^*2*^=0.46[36, 37] for BMI, were used.

**Figure 10.**
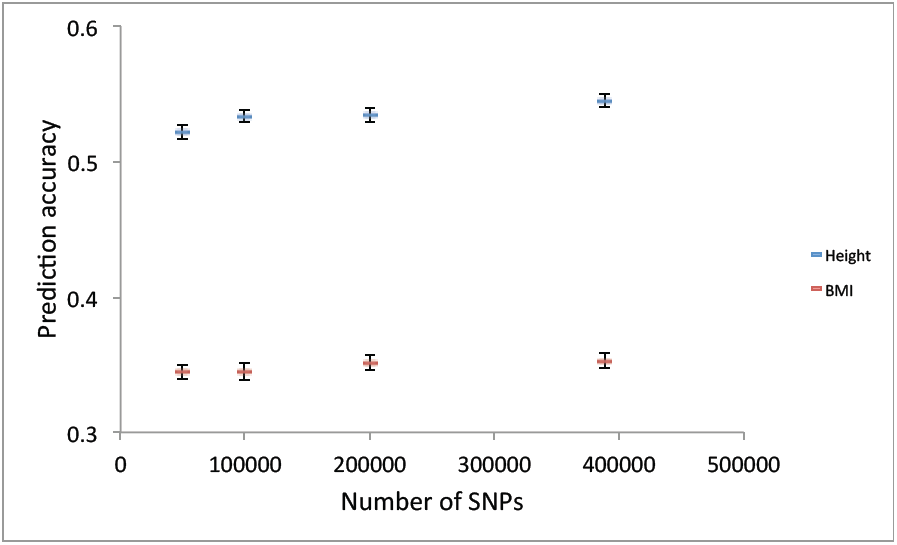
When using Framingham data, the prediction accuracy is not much decreased even with 50,000 SNPs that were randomly selected from 389,265 SNPs.

## DISCUSSION

This work shows a simple approach for modeling genomic prediction in a reference data set that contained several subpopulations that differ in relatedness to the target set, and by modeling these subpopulations as having different effective population size. The model allows assessing the prediction accuracy before actually conducting an experiment so that designing genomic prediction can be precise and effective in animal, plant and human genetics. For example, it can address a question how much the prediction accuracy can be increased by adding 10,000 (conventionally) unrelated individuals into the current experiment consisting of 100 relatives in the reference data. The value for *N*_*e*_ in equation (1) can be approximated based on prior knowledge of a population, and the relatedness of the sample with the target, possibly supported by some genotype information that maybe available on cohorts, or samples thereof. As a reference table, we added approximated value of *N*_*e*_ for structured population of full and half sibs (Supplementary Tables 1 and 2). Prediction in advance indeed relies on arbitrary modelling a number of cohorts, but it would be a useful exercise, as illustrated in the results when considering marker density and various sizes of the subsets of the training data. The theory is also useful for an animal breeder to predict the value of genotyped animals in an own herd versus those in a wider references population consisting of a larger number of more distantly related individuals.

The genotypic and phenotypic information of close and distant relatives of the proband can be effectively used as a part of the unified reference panel that also include a large number of individuals that are not related to the predicted subject to improve the accuracy further as illustrated in Figure 7. For a random sample from the same homogenous population, e.g. within the same breed or ethnicity, an optimal design should consist of both close and distant relatives and unrelated individuals, e.g. a composite design, to maximise the prediction accuracy (Figure 7). That is, the composite design takes advantage of effective information from smaller number of relatives while it also use information from a greater number of unrelated individual.

We showed that the prediction accuracy derived for a population with unrelated individuals turns out to be higher, compared to previous quantifications that overestimated *M*_*e*_ for a larger number of chromosomes[5, 6, 21, 38]. Using the same theory, we also showed that the information from close relatives could increase the accuracy even further, especially for smaller reference populations (Figures 3 – 6). It is important to note that the assumption about using unrelated individuals in estimating empirical *M*_*e*_ from genomic relationship[5] is not strictly necessary and can be relaxed (Eq. (4) and (5)). The theory and empirical observation from simulation study agreed well (Figure 1) even when using a population with a smaller effective population size (*N*_*e*_=50) that consisted of a significant proportion of high relatedness.

Previous studies related to genomic prediction accuracy have suggested that *M*_*e*_ can be derived from the variation in the differences between realized and expected relationships[6, 22], i.e. **D** = **G** – **A** where **G** is a genomic relationship matrix and **A** is a numerator relationship matrix based on pedigree. Those studies validated their results also with simulation. If the individuals used in the training set have a low expected relationship to the target individuals, then there is not much difference between the variations in **D** versus **G**. However, when some closer relatives are used, var(**G**) is larger than var(**D**) and *M*_*e*_ is therefore smaller. Note that non-random sampling of individuals used for the training set can cause a difference between the *N*_*e*_ of the population that was simulated, and the *N*_*e*_ of the data set that was used for prediction.

We have not tested the theory for multi-breed reference populations, i.e. those that are heterogeneous in the sense of consisting of populations from different genetic background, i.e. different breeds or ethnicities, each with different minor allele frequencies, different LD structure and different effects for causal variants. Wientjes et al. (2016)[22] explicitly addressed the problem of different effects for causal variants (i.e. genetic correlation less than one) when combining data from two populations. Individuals from different populations share genomic relationships that are lower than those among members within each population. Evidence in literature suggests low prediction accuracies when using information from different breeds or populations, which could be viewed as predicting from populations with very large *N*_*e*_. Moreover, we have not considered historical population dynamics such as bottleneck and admixture, but assumed a constant *N*_*e*_ over the historical generations, which leads to simplifications that make the formulae tractable and easy to derive. Further work is required to extend the theory accounting for admixture populations and historical population dynamics.

We have shown an improved theory for the prediction of the effective number of chromosome segments, which is a key parameter in genomic prediction accuracy[7]. The theory accounts for the correlation between relationships at different chromosomes and as a result the effective number of chromosome segments is smaller than predicted from previous theory[5, 6, 21]. As a result, the increase of the genomic prediction accuracy appears to be less reliant on higher marker density unless *N*_*e*_ is very large (e.g. > 10,000) (Figure 2), compared to what have been quantified by previous theory[5, 6, 21]. The previous theory overestimates *M*_*e*_ (mostly due to neglecting correlation between chromosomes), therefore underestimates the proportion of genetic variance at QTL captured by markers. Little improvement of prediction accuracy with increasing SNP marker density has been empirically observed in a number of studies[39-41]. This may also have important implication in genomic prediction as to designing marker density in animal, plant and human genetics.

The ability to quantify the accuracy in relation to various degrees of relationships (e.g. close relatives, distant relatives, local or extensive population sample) is important for predicting outcomes of genomic prediction for specific designs. This study has addressed this question, and the theory has been implemented in MTG2 software (https://sites.google.com/site/honglee0707/mtg2). Therefore, a user can know the expected prediction accuracy and the power[26] before designing an experiment of genomic prediction. Our approach can be applied both before and after collecting the data.

## ACKNOWLEDGEMENTS

This research is supported by the Australian National Health and Medical Research Council (APP1080157), the Australian Research Council (DP160102126, FT160100229) and the Australian Sheep Industry Cooperative Research Centre. The Framingham Heart Study is conducted and supported by the National Heart, Lung, and Blood Institute (NHLBI) in collaboration with Boston University (Contract No. N01-HC-25195). This manuscript was not prepared in collaboration with investigators of the Framingham Heart Study and does not necessarily reflect the opinions or views of the Framingham Heart Study, Boston University, or NHLBI. Funding for SHARe Affymetrix genotyping was provided by NHLBI Contract N02-HL-64278. SHARe Illumina genotyping was provided under an agreement between Illumina and Boston University. The authors acknowledge useful discussion with Han Mulder that contributed to a clearer paper.

## DISCLOSURE DECLARATION

The authors declare no competing financial interests.

## References

1. Meuwissen T, Hayes B, Goddard M. Prediction of total genetic value using genome-wide dense marker maps. Genetics. 2001;157(4):1819 – 29.

2. Wray NR, Goddard ME, Visscher PM. Prediction of individual genetic risk to disease from genome-wide association studies. Genome Res. 2007;17:1520–8.

3. Collins FS, Varmus H. A New Initiative on Precision Medicine. New England Journal of Medicine. 2015;372(9):793–5. doi:10.1056/NEJMp1500523. PubMed PMID: 25635347.

4. Jannink J-L, Lorenz AJ, Iwata H. Genomic selection in plant breeding: from theory to practice. Brief Funct Genomics. 2010;9(2):166–77. doi: 10.1093/bfgp/elq001. 5.

5. Goddard ME, Hayes BJ, Meuwissen THE. Using the genomic relationship matrix to predict the accuracy of genomic selection. Journal of Animal Breeding and Genetics. 2011;128(6):409–21. doi: 10.1111/j.1439-0388.2011.00964.x.

6. Goddard ME. Genomic selection: prediction of accuracy and maximisation of long term response. Genetica. 2009;136:245–57. doi: 10.1007/s10709-008-9308-0.

7. Lee SH, Weerasinghe WMSP, Wray N, Goddard M, Van der Werf J. Using information of relatives in genomic prediction to apply effective stratified medicine. Scientific Reports. 2017;7:42091. doi: DOI: 10.1038/srep42091.

8. Tucker G, Loh P-R, MacLeod Iona M, Hayes Ben J, Goddard Michael E, Berger B, et al. Two-Variance-Component Model Improves Genetic Prediction in Family Datasets. The American Journal of Human Genetics. 2015;97(5):677–90. doi: 10.1016/j.ajhg.2015.10.002.

9. de los Campos G, Vazquez AI, Fernando R, Klimentidis YC, Sorensen D. Prediction of Complex Human Traits Using the Genomic Best Linear Unbiased Predictor. PLoS Genet. 2013;9(7):e1003608. doi: 10.1371/journal.pgen.1003608.

10. Makowsky R, Pajewski NM, Klimentidis YC, Vazquez AI, Duarte CW, Allison DB, et al. Beyond Missing Heritability: Prediction of Complex Traits. PLoS Genet. 2011;7(4):e1002051. doi: 10.1371/journal.pgen.1002051.

11. Aulchenko YS, Struchalin MV, Belonogova NM, Axenovich TI, Weedon MN, Hofman A, et al. Predicting human height by Victorian and genomic methods. Eur J Hum Genet. 2009;17(8):1070–5. Epub 2009/02/19. doi: 10.1038/ejhg.2009.5. PubMed PMID: 19223933; PubMed Central PMCID: PMCPmc2986552.

12. Lee S, van der Werf J, Hayes B, Goddard M, Visscher P. Predicting unobserved phenotypes for complex traits from whole-genome SNP data. PLoS Genet. 2008;4(10):e1000231. doi: 10.1371/journal.pgen.1000231. PubMed PMID: WOS:000261480900026.

13. Clark SA, Hickey JM, Daetwyler HD, van der Werf JH. The importance of information on relatives for the prediction of genomic breeding values and the implications for the makeup of reference data sets in livestock breeding schemes. Genet Sel Evol. 2012;44:4. Epub 2012/02/11. doi: 10.1186/1297-9686-44-4. PubMed PMID: 22321529; PubMed Central PMCID: PMCPmc3299588.

14. Legarra A, Robert-Granie C, Manfredi E, Elsen JM. Performance of genomic selection in mice. Genetics. 2008;180(1):611–8. Epub 2008/09/02. doi: 10.1534/genetics.108.088575. PubMed PMID: 18757934; PubMed Central PMCID: PMCPmc2535710.

15. Habier D, Fernando RL, Garrick DJ. Genomic BLUP Decoded: A Look into the Black Box of Genomic Prediction. Genetics. 2013;194(3):597–607. doi: 10.1534/genetics.113.152207.

16. Pszczola M, Strabel T, Mulder HA, Calus MP. Reliability of direct genomic values for animals with different relationships within and to the reference population. J Dairy Sci. 2012;95(1):389–400. Epub 2011/12/24. doi: 10.3168/jds.2011-4338. PubMed PMID: 22192218.

17. Hayes B, Visscher P, Goddard M. Increased accuracy of artificial selection by using the realized relationship matrix. Genet Res. 2009;91:47 - 60. PubMed PMID: doi:10.1017/S0016672308009981.

18. van der Werf JHJ, Clark SA, Lee SH, editors. Predicting genomic selection accuracy from heterogeneous sources. Association for the Advancement of Animal Breeding and Genetics Conference; 2015; Lorne, Australia: AAABG.

19. Wientjes YC, Veerkamp RF, Calus MP. The effect of linkage disequilibrium and family relationships on the reliability of genomic prediction. Genetics. 2013;193(2):621–31. Epub 2012/12/26. doi: 10.1534/genetics.112.146290. PubMed PMID: 23267052; PubMed Central PMCID: PMCPMC3567749.

20. Rabier C-E, Barre P, Asp T, Charmet G, Mangin B. On the Accuracy of Genomic Selection. PLoS ONE. 2016;11(6):e0156086. doi: 10.1371/journal.pone.0156086.

21. Meuwissen T, Hayes B, Goddard M. Accelerating Improvement of Livestock with Genomic Selection. Annual Review of Animal Biosciences. 2013;1(1):221–37. doi:10.1146/annurev-animal-031412-103705. PubMed PMID: 25387018.

22. Wientjes YCJ, Bijma P, Veerkamp RF, Calus MPL. An Equation to Predict the Accuracy of Genomic Values by Combining Data from Multiple Traits, Populations, or Environments. Genetics. 2016;202:799–823. doi: 10.1534/genetics.115.183269.

23. Wray NR, Lee SH, Mehta D, Vinkhuyzen AAE, Dudbridge F, Middeldorp CM. Research Review: Polygenic methods and their application to psychiatric traits. Journal of Child Psychology and Psychiatry. 2014;55(10):1068–87. doi: 10.1111/jcpp.12295.

24. Daetwyler HD, Villanueva B, Woolliams JA. Accuracy of predicting the genetic risk of disease using a genome-wide approach. PLoS ONE. 2008;3(10):e3395.

25. Sved JA. Linkage disequilibrium and homozygosity of chromosome segments in finite populations. Theor Popul Biol. 1971;2(2):125–41. PubMed PMID: 5170716.

26. Lee SH, Wray NR. Novel genetic analysis for case-control genome-wide association studies: quantification of power and genomic prediction accuracy. PLoS ONE. 2013;8(8):e71494. doi: 10.1371/journal.pone.0071494. PubMed PMID: WOS:000323425700059.

27. Lee SH, Wray NR, Goddard ME, Visscher PM. Estimating missing heritability for disease from genome-wide association studies. Am J Hum Genet. 2011;88:294–305. PubMed PMID: s0002-9297(11)00020-6 DOI - 10.1016/j.ajhg.2011.02.002.

28. Chow S, Sahao J, Wang H. Sample Size Calculations in Clinical Research. 2nd Ed ed 2008.

29. MacCluer JW, VandeBerg JL, Read B, Ryder OA. Pedigree analysis by computer simulation. Zoo Biology. 1986;5:147–60.

30. Lee S, van der Werf J. The efficiency of designs for fine-mapping of quantitative trait loci using combined linkage disequilibrium and linkage. Genetics Selection Evolution. 2004;36(2):145–61. doi: 10.1051/gse:2003056. PubMed PMID: WOS:000188936300001.

31. Roach JC, Glusman G, Smit AFA, Huff CD, Hubley R, Shannon PT, et al. Analysis of Genetic Inheritance in a Family Quartet by Whole-Genome Sequencing. Science. 2010;328(5978):636–9. doi: 10.1126/science.1186802.

32. Splansky GL, Corey D, Yang Q, Atwood LD, Cupples LA, Benjamin EJ, et al. The Third Generation Cohort of the National Heart, Lung, and Blood Institute's Framingham Heart Study: design, recruitment, and initial examination. Am J Epidemiol. 2007;165(11):1328–35. Epub 2007/03/21. doi: 10.1093/aje/kwm021. PubMed PMID: 17372189.

33. Silventoinen K, Sammalisto S, Perola M, Boomsma DI, Cornes BK, Davis C, et al. Heritability of adult body height: a comparative study of twin cohorts in eight countries. Twin research : the official journal of the International Society for Twin Studies. 2003;6(5):399–408. Epub 2003/11/20. doi: 10.1375/136905203770326402. PubMed PMID: 14624724.

34. Macgregor S, Cornes BK, Martin NG, Visscher PM. Bias, precision and heritability of self-reported and clinically measured height in Australian twins. Hum Genet. 2006;120(4):571–80. Epub 2006/08/26. doi: 10.1007/s00439-006-0240-z. PubMed PMID: 16933140.

35. Visscher PM, Medland SE, Ferreira MAR, Morley KI, Zhu G, Cornes BK, et al. Assumption-free estimation of heritability from genome-wide identity-by-descent sharing between full siblings. PLoS Genet. 2006;2(3):e41.

36. Elks CE, den Hoed M, Zhao JH, Sharp SJ, Wareham NJ, Loos RJ, et al. Variability in the heritability of body mass index: a systematic review and meta-regression. Frontiers in endocrinology. 2012;3:29. Epub 2012/05/31. doi: 10.3389/fendo.2012.00029. PubMed PMID: 22645519; PubMed Central PMCID: PMCPMC3355836.

37. Wilson JG, Rotimi CN, Ekunwe L, Royal CD, Crump ME, Wyatt SB, et al. Study design for genetic analysis in the Jackson Heart Study. Ethnicity & disease. 2005;15(4 Suppl 6):s6-30-7. Epub 2005/12/02. PubMed PMID: 16317983.

38. Goddard ME, Hayes BJ. Mapping genes for complex traits in domestic animals and their use in breeding programmes. Nat Rev Genet. 2009;10(6):381–91.

39. Su G, Brøndum RF, Ma P, Guldbrandtsen B, Aamand GP, Lund MS. Comparison of genomic predictions using medium-density (∼54,000) and high-density (∼777,000) single nucleotide polymorphism marker panels in Nordic Holstein and Red Dairy Cattle populations. Journal of Dairy Science. 2012;95(8):4657–65. doi: http://dx.doi.org/10.3168/jds.2012-5379.

40. VanRaden, Paul M., O'Connell, Jeffrey R., Wiggans GR, Weigel KA. Genomic evaluations with many more genotypes. Genetics Selection Evolution. 2011;43(1):1–11. doi: 10.1186/1297-9686-43-10.

41. Moghaddar N, Swan AA, van der Werf JHJ. Accuracy of genomic prediction for Merino wool traits using high-density marker genotypes. Proc Assoc Advmt Breed Genet 2015;21:165–8.

